# Last-century vegetation dynamics in a highland Pyrenean national park and implications for conservation

**DOI:** 10.1101/2024.03.01.582940

**Authors:** V. Rull, A. Blasco, J. Sigró, T. Vegas-Vilarrúbia

## Abstract

Ecological records from before and after the creation of natural parks are valuable for informing conservation and restoration actions. Such records are often unavailable, but high-resolution paleoecological studies may provide useful information. This paper presents a sub-decadal paleoecological reconstruction of vegetation and landscape in a national park in the Pyrenean highlands, established in the 1950s. The park lands were traditionally been used for small-scale cultivation, extensive grazing, forest exploitation and, since 1910, hydroelectricity generation following the damming of numerous glacial lakes. A significant finding is that present-like forests, with negligible changes in composition, have dominated the landscape during the study period. Major vegetation changes involved shifts in forest cover, influenced by both climatic and anthropic factors. Interestingly, the creation of the park in 1955 and the initial restrictions on forest exploitation in 1975 did not significantly affect vegetation cover or composition. Forest expansion did not occur significantly until the 1980s when the park was enlarged, and forest exploitation was further restricted. This expansion peaked in the mid-1990s coinciding with a warming trend and a decrease in fire incidence, before declining due to warmer and drier climates. This decline in forest cover occurred concurrently with the ongoing global forest dieback phenomenon and may be exacerbated by the predicted global warming in this century, which could also increase fire incidence due to the accumulation of dead wood. Under current conservation measures, the main threats are global warming, fire and, on a more local scale, the massification of tourism. Expanding the park and implementing forest restoration actions on degraded terrains surrounding the park could only be beneficial.

## 1. Introduction

Understanding the dynamics of vegetation and landscape over the last centuries, influenced by both climate change and human management, is crucial for optimizing conservation and restoration efforts. However, long-term ecological studies spanning 50 years or more are exceedingly rare, which highlights the value of high-resolution paleoecological records derived from recent sediments (Willis & Birks, 2006; Willis et al., 2007, 2011; Rull & Vegas-Vilarrúbia, 2011; Vegas-Vilarrúbia et al., 2011; Davies et al., 2014; Rull, 2014; Seddon et al., 2014). Natural parks and other areas designated for protection in the 20th century are particularly well-suited for these studies. The recent sediments in their wetlands can provide detailed accounts of ecological changes following the introduction of conservation measures. These insights are useful for shaping management policies, offering precise tools for improved environmental stewardship of each site. Indeed, site-specific studies have proven to be highly beneficial, as they leverage paleoecological insights to inform conservation strategies by shedding light on ecosystem dynamics, ecological responses, and critical thresholds (Davies et al., 2014).

In the Iberian Pyrenees, there are two highland national parks—Aigüestortes i Estany de Sant Maurici (PNAESM) and Ordesa y Monte Perdido (PNOMP)—alongside other protected sites (natural parks), which boast numerous lakes and peat bogs containing records spanning from the Lateglacial to the present (Rull & Vegas-Vilarrúbia, 2021). However, high-resolution paleoecological records from the last century are scarce. The most detailed 20th-century Pyrenean record, featuring an average of approximately six years per sampling interval, was obtained from the annually-laminated (varved) sediments of the mid-elevation Lake Montcortès (1027 m) (Rull et al., 2021). Other records covering the last centuries typically have resolutions at centennial or multicentennial time scales (review in Rull & Vegas-Vilarrúbia, 2022). Therefore, no high-resolution paleoecological records of the last century that may inform conservation are available for the PNAESM or the PNOMP.

This study focuses on the PNAESM, established in the mid-20th century. The specific target is Lake Sant Maurici, the park’s most iconic lake, from which a sediment core representing approximately the last century was obtained and analyzed palynologically at sub-decadal resolution (averaging 3.5 years per sampling interval). The main objectives of this research were: 1) to reconstruct the shifts in vegetation and landscape, with a particular emphasis on forest dynamics and fire incidence; 2) to correlate these trends with available climatic data and documented human activities, especially forest exploitation, grazing, and hydroelectricity generation; 3) to evaluate the impact of lake damming and the establishment of the park on the dynamics of vegetation and landscape; 4) to assess the resilience of highland forests against both natural and anthropogenic ecological pressures; 5) to utilize the findings to inform conservation practices aimed at enhancing governance and management of the park. This high-resolution paleoecological reconstruction offers a unique record of vegetation dynamics and associated ecological drivers, unavailable through modern ecological studies. Additionally, this record falls within a temporal window that facilitates the integration of modern ecological and paleoecological data, thus paving the way for the creation of continuous and truly long-term ecological datasets (Rull, 2014). Such records are more informative for conservation actions than short-term ecological studies.

## 2. Study area

### 2.1. General description

The PNAESM (Fig. 1) was created in 1955, initially covering an area of 9,851 ha. Before this, the lands were privately owned and the main activities were subsistence agriculture, extensive grazing, usually transhumant, hunting, fishing and gathering. By 1975, forest exploitation had been restricted, but hydroelectricity generation—which began in 1910 and reached its peak from 1946 to 1960—and extensive grazing continued. In some sectors of the park, however, forest exploitation continued, due to previous agreements with private owners (Gil-Farrero, 2021, 2022). Hydroelectric power was generated by damming and underground drilling of a number of the abundant (>270) alpine lakes of glacial origin, situated at elevations between approximately 2000 and 2500 m (Catalan et al., 1997). However, the hydroelectric plants are located outside the park’s boundaries. Sant Maurici, one of the lakes impacted, was dammed in 1953, two years prior to the park establishment. Between 1985 and 1996, the park’s lands were reclassified as public and its area expanded to its current size of 14,119 ha, with an additional peripheral protection zone of 26,733 ha (Fig. 1). Under this new administrative regime, the exploitation of natural resources, except for hydroelectricity and animal husbandry, became more restricted, particularly forest exploitation. Since the creation of the park, conflicts among the different actors involved – especially political/administrative sectors and private/communal owners – have been continuous due to the different interests and perceptions on how nature and natural resources should be managed (Gil-Farrero, 2021, 2022).

**Figure 1.**
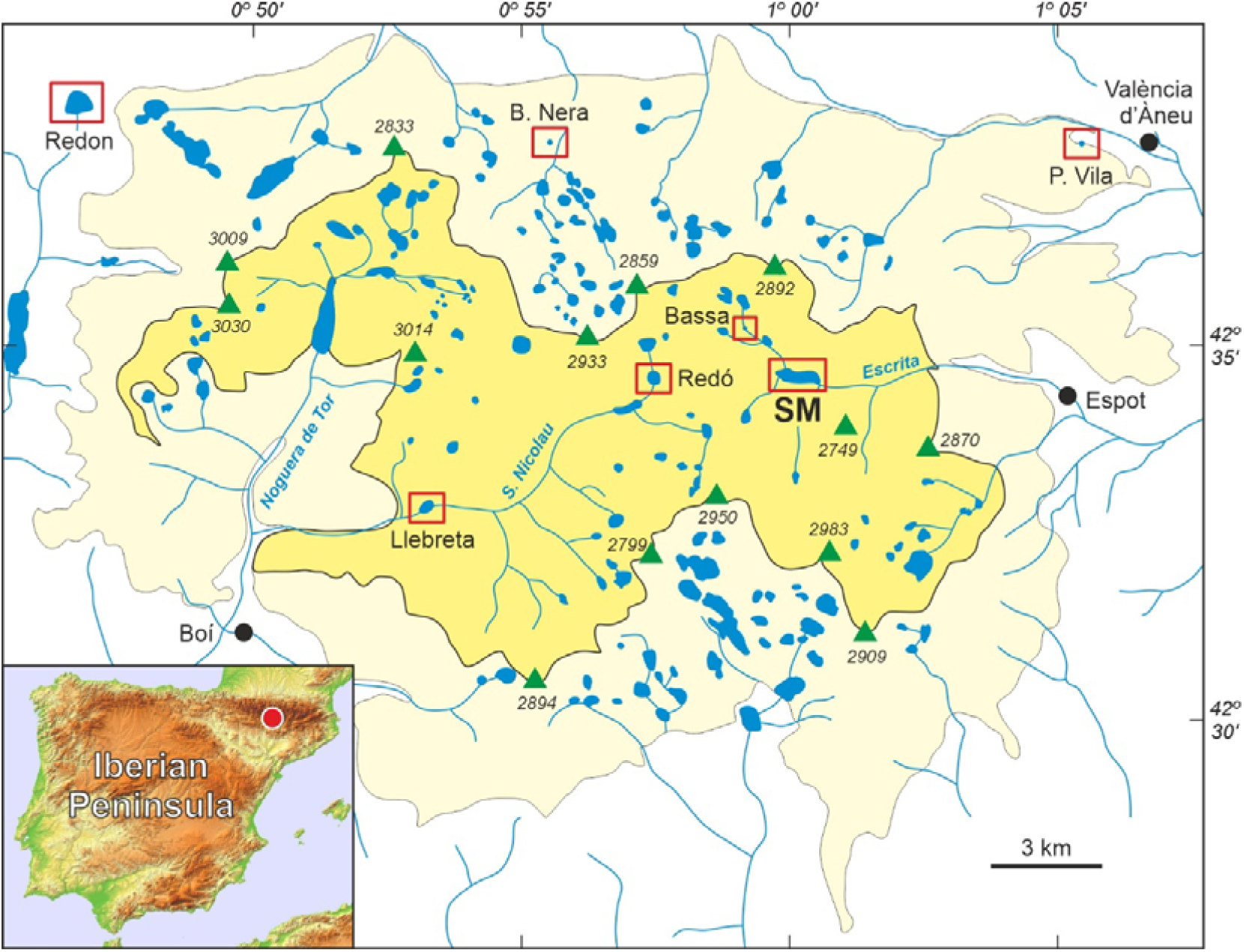
Sketch map of the Pyrenean “Aigüestortes i Estany de Sant Maurici” National Park (PNAESM) and its location in the Iberian Peninsula (red dot). The park itself is in yellow and the peripheral protection zone is in light yellow. Green triangles represent the highest peaks (elevations in m a.s.l), and lakes and rivers are in blue. The localities (lakes, peat bogs) with Lateglacial/Holocene pollen records are highlighted by red boxes (SM, Lake Sant Maurici). Redrawn and modified from Rull et al. (2021).

Topographically, the PNAESM ranges from approximately 1600 m (1,300 m including the peripheral buffer zone) to the highest peaks, which rise slightly above 3000 m (Carrillo & Aniz, 2013). Geologically, the park predominantly consists of Paleozoic formations, including Ordovician, Silurian, Devonian, and Carboniferous shales, interspersed with abundant Carboniferous granitic intrusions. The current landscape has been primarily sculpted by glacial activity during the Pleistocene. This glacial action has forged a distinctive glacial landscape characterized by features such as horns, ridges, cirques, U-shaped valleys, “roches moutonées”, moraines and lakes, all superimposed on the Paleozoic bedrock (Rodríguez-Fernández, 2010).

According to the record from the AEMET-9660 meteorological station, located near the lake at an elevation of 1920 m, the average annual temperature is 5.4°C, and the average annual precipitation is 1058 mm for the period 1988–2017. The highest average monthly temperatures, slightly above 14°C, occur in summer (July–August), while the lowest, approximately −1.5°C, are observed in winter (January–February). In terms of precipitation, monthly averages consistently exceed 60 mm, peaking in May (188 mm) and November (110 mm). The lowest precipitation levels were noted in February (63 mm) and July (65 mm) (Rull et al., 2021). Additionally, an integrated instrumental climatic record for the entire PNAESM during the 20th century has been compiled recently (Sigró et al., 2022), covering a substantial portion of the timeframe addressed in this study.

The general distribution of vegetation is primarily governed by marked elevational gradients in climatic parameters, especially temperature, which decreases at an average rate of 0.6°C per 100 m in elevation (Carrillo & Aniz, 2013). The montane belt, ranging from 1200 to 1600/1800 m and predominantly found in the peripheral zone, features deciduous and mixed forests with species such as *Abies alba* (silver fir), *Fagus sylvatica* (beech), *Corylus avellana* (hazel), *Betula pendula* (silver birch), *Fraxinus excelsior* (ash), and *Pinus sylvestris* (Scots pine). The subalpine belt, spanning 1600 to 2300/2500 m, is mainly comprised of conifer forests with *Abies alba* and *Pinus mugo* subsp. *uncinata* (black pine, also known as *P. uncinata*), with the upper boundary of this belt marking the treeline, which is located between 2300 and 2500 m elevation, varying with local slope characteristics. Above this treeline lies the alpine belt, dominated by Festuca eskia grasslands. The alpine wetlands, encircling lakes and bogs, are dominated by species such as *Carex nigra, Eriophorum angustifolium*, and *Scirpus cespitosus*. Beyond 2700 m, in the subnival zone, cushion-like plants, including *Saxifraga* spp. and *Silene acaulis*, can be found thriving in rocky areas (Carrillo & Aniz, 2013).

Lake Sant Maurici (42°34⍰53⍰N, 1°00⍰12⍰E; 1914 m elevation) (Fig. 1) is located in a glacial cirque within the subalpine belt, surrounded by the typical high-altitude conifer forests of black pine and birch. Originally, the lake was much smaller and shallower than it is today. In 1953 CE, the construction of a 16-m high dam transformed this small pond into a reservoir approximately 1 km long and 200 m wide (Fig. 2), with a maximum depth of 25 m. This reservoir is now used for hydroelectricity generation, irrigation, and human consumption (Catalan et al., 1997). The damming and related constructions led to the removal of vegetation around the dam and the introduction of ruderal species, increased soil erosion around the access routes, and flooded the surrounding terrains. This flooding irreversibly destroyed the basin floor’s vegetation not adapted to inundation and water-level fluctuations, primarily affecting conifer forests and peat bogs along the lake margins, which were habitats for many unique floristic elements of the Pyrenean highlands (Vigo, 2008). The aquatic ecosystems also suffered significantly from the raised water levels, eutrophication, and oxygen depletion due to the decomposition of dead plant biomass (Catalan et al., 1997).

**Figure 2.**
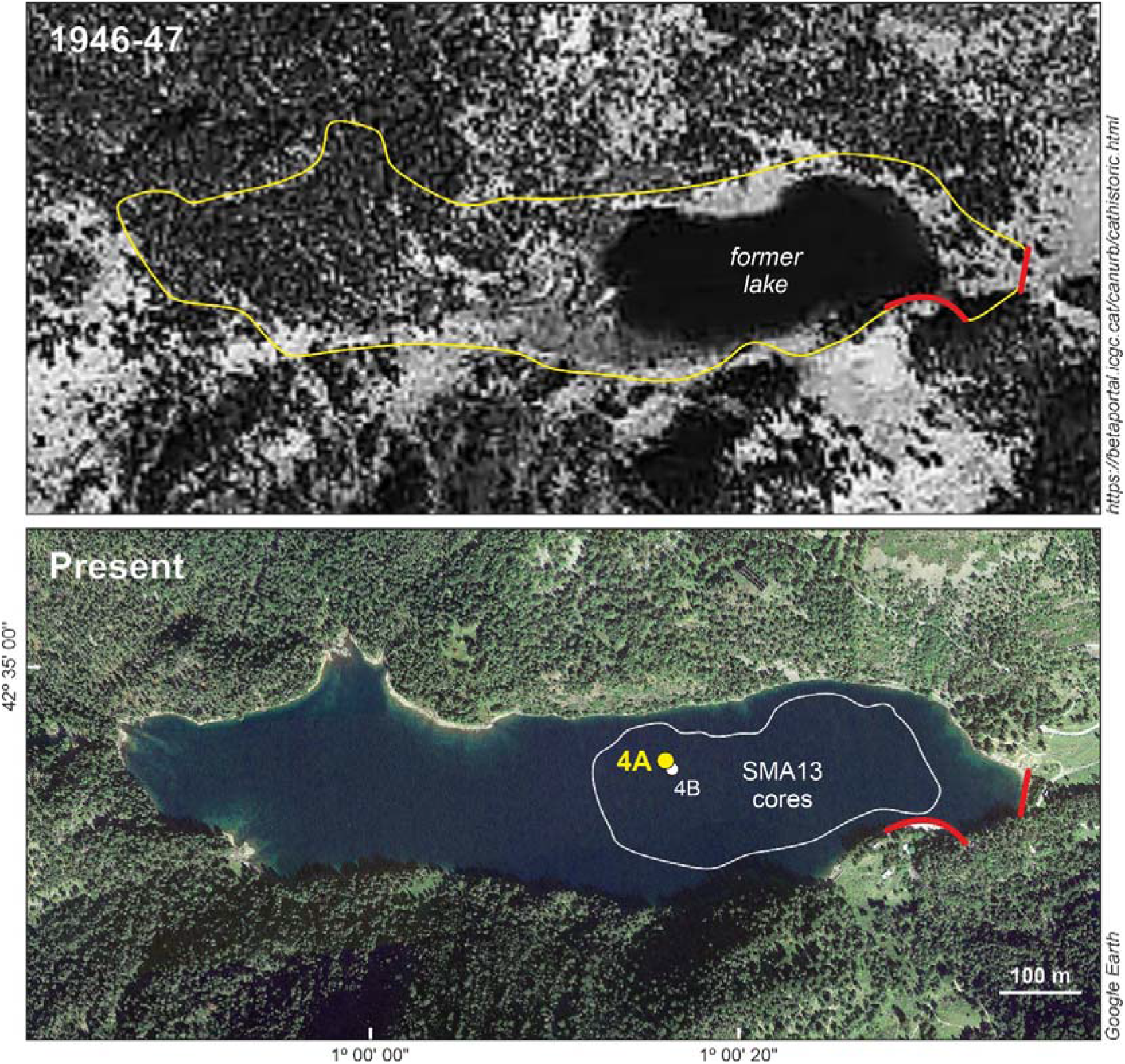
Lake Sant Maurici before and after damming (1953 CE). The upper panel shows the former glacial lake and the contour of the present reservoir (yellow line), whereas the lower panel displays the present reservoir and the contour of the former lake (white line). Damming areas are indicated by red lines. The core used in this study (SMA13-4A) is highlighted in yellow and the core studied by Calero et al. (2016) and Rull et al. (2021) (SMA13-4B) is indicated in white. See Calero et al. (2022) and Rull (2023) for more details.

### 2.2. Previous studies

Palynological studies conducted in the PNAESM and its surroundings (Fig. 1) have reconstructed the Lateglacial and Holocene vegetation dynamics with multidecadal to centennial resolution (Esteban, 2003; Catalan et al., 2013; Rodríguez et al., 2021). In Sant Maurici, a 9 m-deep core (SMA13-4B) was retrieved from the edge of the former glacial lake (Fig. 2). This core contains a fragmentary Holocene record, due to sedimentation interruptions caused by water-level fluctuations (Calero et al., 2016). The Late Holocene section of this core encompasses the Bronze Age (4.3-3.5 cal kyr BP) and the Middle Ages (1.5-0.3 cal kyr BP), with a significant sedimentary gap omitting the interval corresponding to the Iron Age and the Roman period (3.5-1.5 cal kyr BP). The analysis revealed that the highland forests around the lake maintained stability in abundance and composition, even through temperature and moisture shifts associated with the Bronze Age Warm Period (BAWP), the Dark Ages Cold Period (DACP), the Medieval Climate Anomaly (MCA), and the Little Ice Age (LIA). Human activities in the area were intermittent and of low intensity, exerting minimal impact on the forest dynamics (Rull et al., 2021).

Therefore, unlike most highland Pyrenean landscapes, which experienced extensive deforestation and anthropogenic transformation from the Bronze Age to the Middle Ages, the forests around Sant Maurici demonstrated remarkable constancy over time. It has been suggested that Sant Maurici may have served as a microrefugium for highland forests during the Late Holocene (Rull & Vegas-Vilarrúbia, 2021). A 1 m-deep gravity core, retrieved from the same location (SMA13-4B-1G), provided a record spanning the last 0.7 cal kyr BP. This record includes a significant gap between 0.6 cal kyr BP and the present, but offers a complete record of the last century in the upper 22 cm. However, high-resolution pollen analysis of this segment was hindered by the scarcity of material remaining after sampling for ^210^Pb dating; only four samples were analyzed, which proved insufficient for a detailed reconstruction of vegetation and landscape dynamics during the 20th century (Camarero et al., 2013).

## 3. Methods

### 3.1. Coring and dating

The Sant Maurici core used in this study (SMA13-4A; 141 cm depth) was collected in October 2013 using a UWITEC gravity corer with a diameter of 63 cm, at a water depth of approximately 18 m, in the vicinity of the former glacial lake, near the previously mentioned parallel core SMA13-4B (Fig. 2). The dating was conducted at the Department of Physics of the Autonomous University of Barcelona. ^210^Pb was analyzed under the assumption of secular equilibrium with its daughter ^210^Po. Following microwave-assisted acid digestion, ^210^Po was quantified using alpha spectrometry (Sanchez-Cabeza et al., 1998). To calculate the sedimentation rate, the constant rate of supply (CRS) model for ^210^Pb was applied (Appleby & Oldfield, 1978). Supported ^210^Pb levels were determined from a composite of three deep samples using gamma spectrometry, specifically through the measurement of ^214^Pb emission lines at 295 keV and 352 keV, assuming secular equilibrium with ^226^Ra.

### 3.2. Pollen analysis

For pollen analysis, 25 samples of 2 cm^3^ were collected at regular intervals using a volumetric syringe. These samples underwent digestion with NaOH for disaggregation, HCl for carbonate removal and HF for silicate removal, followed by centrifugation with Thoulet solution (density 2 g/cm^3^) for mineral separation (Bennett & Willis, 2001). Prior to processing, two tablets of *Lycopodium* spores (Batch 280521 291; 13,761±448 spores/tablet) were added to each sample (Stockmarr, 1971). The processing took place at the Archaeobotany Lab of the Catalan Institute of Human Paleoecology and Social Evolution (IPHES). Pollen and spore identification utilized Moore et al. (1991), Reille (1992), and the author’s (VR) own reference collection, which comprises over 700 species/subspecies from the Iberian flora. Non-pollen palynomorphs (NPP) identification followed López-Vila et al. (2014) and the NPP Image Database (accessible at https://non-pollen-palynomorphs.uni-goettingen.de/, last visited 3 May 2023). While fungal spores were generally not identified, those from coprophilous fungi, indicating grazing and hence human impact, received special attention (Cugny et al., 2010; Baker et al., 2013; Gauthier & Jouffroy-Bapicot, 2021; Lee et al., 2022). Spores of *Glomus*, a mycorrhizal fungus indicative of soil erosion following local forest clearance in the study area, were also identified (López-Vila et al., 2014). Charcoal particles served as proxies for the fire regime and were categorized into two main size groups to differentiate between regional fires (<150 μm) and fires occurring near the coring site (>150 μm) (Whitlock & Larsen, 2001).

Counting was conducted until reaching a minimum pollen sum of 300 grains, along with the stabilization of confidence intervals and saturation of diversity (Rull, 1987). In several samples, *Pinus* pollen was overwhelmingly abundant, with counts halted at 100-150 grains. Beyond this point, counts were extrapolated using *Lycopodium* spores. For these samples, the specified statistical criteria were applied to the non-*Pinus* pollen types. The pollen sum excluded aquatic and semi-aquatic plants (Cyperaceae, *Epilobium/Oenothera, Ranunculus*) to focus on terrestrial vegetation. Consistent with standard practices in pollen analysis, percentages were calculated to deduce vegetation composition, while pollen accumulation rates (PAR) served as proxies for plant cover (Theuerkauf & Couwenberg, 2018).

## 4. Results and interpretation

### 4.1. Chronology and sedimentation

The results of the ^210^Pb dating are presented in Table 1. Samples with dating errors exceeding the sampling intervals (more than ±6 years) were excluded from this study. Consequently, the period analyzed spans 88 years, from 1919 to 2007 CE, with an average resolution of 3.5 years. While sample 20P was utilized for pollen analysis, it was not used for dating purposes and its age was interpolated. The age-depth model is illustrated alongside the pollen diagram in Figure 3 to enhance visualization. Sediment accumulation rates averaged 0.285 cm per year (cm/y), displaying an increasing trend that peaked at 0.463 cm/y around the mid-section (11.2 cm depth, corresponding to 1974 CE) before declining. To aid in comparison, the dates of significant human activities discussed in section 2.1 are marked on the chronological scale derived from the age-depth model, as shown in Fig. 3.

**Table 1.** Results of ^210^ Pb dating using the CRS model and the depth of sample tops. SAR, sediment accumulation rates.

| Sample | Depth (cm) | Excess $^{210}\text{Pb}$ (Bq kg$^{-1}$) | Date (year) | SAR (cm y$^{-1}$) |
| --- | --- | --- | --- | --- |
| 1P | 1.0 | 292.5±11.1 | 2007±1 | 0.282±0.013 |
| 2P | 2.0 | | 2005±1 | 0.301±0.017 |
| 3P | 3.1 | 360.4±21.5 | 2002±1 | 0.212±0.015 |
| 4P | 4.1 | | 1997±1 | 0.269±0.017 |
| 5P | 5.1 | 228.4±11.1 | 1993±1 | 0.291±0.017 |
| 6P | 6.1 | | 1990±1 | 0.266±0.021 |
| 7P | 7.1 | 164.9±16.0 | 1986±1 | 0.343±0.037 |
| 8P | 8.1 | | 1983±1 | 0.344±0.040 |
| 9P | 9.2 | 98.3±11.5 | 1980±1 | 0.384±0.049 |
| 10P | 10.2 | | 1977±1 | 0.322±0.043 |
| 11P | 11.2 | 76.4±9.4 | 1974±1 | 0.463±0.062 |
| 12P | 12.2 | | 1972±2 | 0.394±0.051 |
| 13P | 13.2 | 85.8±9.2 | 1969±2 | 0.303±0.037 |
| 14P | 14.3 | | 1966±2 | 0.298±0.033 |
| 15P | 15.3 | 91.1±6.9 | 1963±2 | 0.291±0.029 |
| 16P | 16.3 | | 1959±2 | 0.307±0.038 |
| 17P | 17.3 | 77.3±9.9 | 1956±2 | 0.341±0.052 |
| 18P | 18.3 | | 1953±2 | 0.333±0.052 |
| 19P | 19.9 | 52.5±6.9 | 1950±3 | 0.256±0.043 |
| 20P<sup>1</sup> | 20.4 | | 1947±3 | 0.248±0.042 |
| 21P | 21.4 | 60.5±7.6 | 1944±3 | 0.239±0.041 |
| 22P | 22.4 | | 1937±4 | 0.199±0.037 |
| 23P | 23.4 | 61.8±8.2 | 1932±4 | 0.146±0.030 |
| 24P | 24.4 | | 1925±5 | 0.161±0.038 |
| 25P | 25.5 | 43.5±7.7 | 1919±6 | 0.120±0.035 |
| 1C<sup>2</sup> | 27.0 | | 1911±8 | 0.070±0.039 |
| 2C<sup>2</sup> | 28.0 | 34.8±17.7 | 1889±11 | 0.043±0.037 |
| 3C<sup>2</sup> | 29.0 | | 1865±15 | 0.031±0.031 |
| 4C<sup>2</sup> | 30.0 | | 1832±32 | ND |
| 5C<sup>2</sup> | 31.1 | <0 | ND | ND |

<sup>1</sup>Only for pollen analysis (interpolated age)
<sup>2</sup>Not used in this work

**Figure 3.**
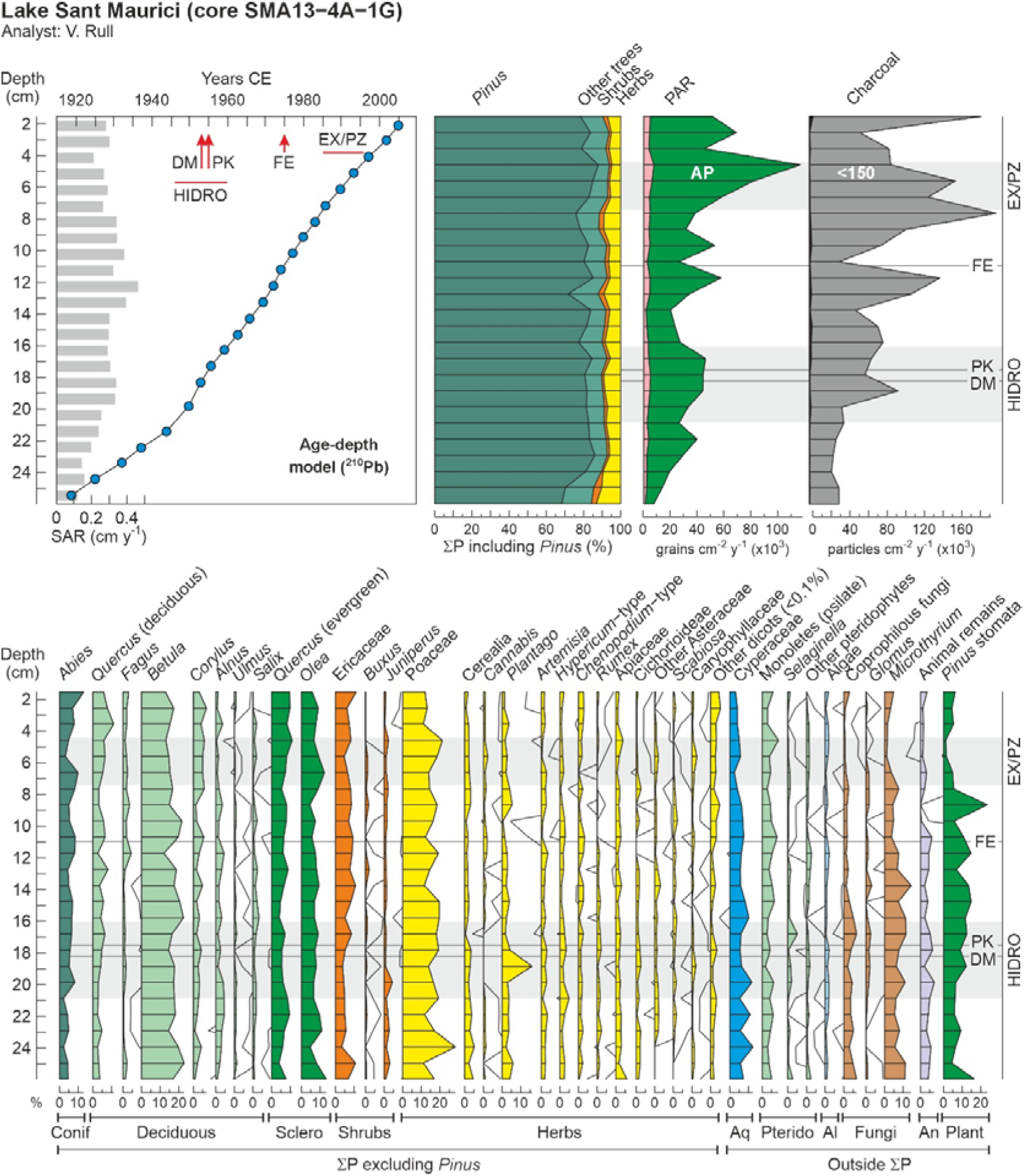
Age-depth model and pollen diagrams for the Sant Maurici core SMA13-4A-1G. The upper panel shows the age-depth model (Table 1), the summary percentage diagram using the main pollen sum, which includes *Pinus*, the pollen accumulation rates (PAR) of arboreal (green) and non-arboreal pollen (pink), and the charcoal influx (particles larger than 150 μm in black and the others in gray). Abbreviations: HIDRO, maximum development of hydroelectrical exploitation; DM, damming of Lake Sant Maurici; PK, creation of the PNAESM; FE, restriction of forest exploitation; EX/PZ, park expansion and creation of the protection zone. The lower panel displays the abundance of palynomorphs other than *Pinus*, with respect to the reduced pollen sum (excluding *Pinus*), using the pollen types above 01% of the total. Other dicots (<0.1%) include *Castanea, Juglans, Tilia, Acer, Ilex, Ephedra, Helianthemum, Galium, Centaurea, Ribes, Valerianella, Geranium, Malvaceae, Lupinus, Phyteuma, Filipendula, Urtica, Drosera, Epilobium/Oenothera and Ranunculus*. Other pteridophytes include *Asplenium, Thelypteris, Isoetes, Polypodium, Sphagnum, Cryptogramma and psilate triletes*. Algae include *Pediastrum, Spirogyra, Staurastrum, Bulbochaete, Botryococcus, Mougeotia* and *Debarya*. Coprophilous fungi are *Sporormiella, Sordaria* and *Podospora*. Animal remains include chironomids, tardigrada *(Macrobiotus)*, rotifers *(Habrotrocha)*, acari, cladocera, tecamoebae *(Arcella)* and platelminthes *(Neorhabdocoela)* (Fig. 4).

### 4.2. Vegetation dynamics

In total, 90 palynomorph types were identified and counted: 56 types were pollen, 9 were pteridophyte spores, and 25 were NPP (8 algae, 5 fungi, 7 animals, 1 plant and 4 of unknown origin). The most abundant NPP types are depicted in Fig. 4 to support future research in the area, where studies on NPP are scarce. The stratigraphic distribution of these palynomorphs is illustrated in the integrated pollen diagram (Fig. 3). *Pinus* pollen was overwhelmingly dominant, showing minimal variation (accounting for 81±10% of the main pollen sum, which averaged 1274 grains). To mitigate potential masking effects on the visibility of other pollen types, a reduced pollen sum was calculated by excluding *Pinus*, resulting in an average of 233 grains.

**Figure 4.**
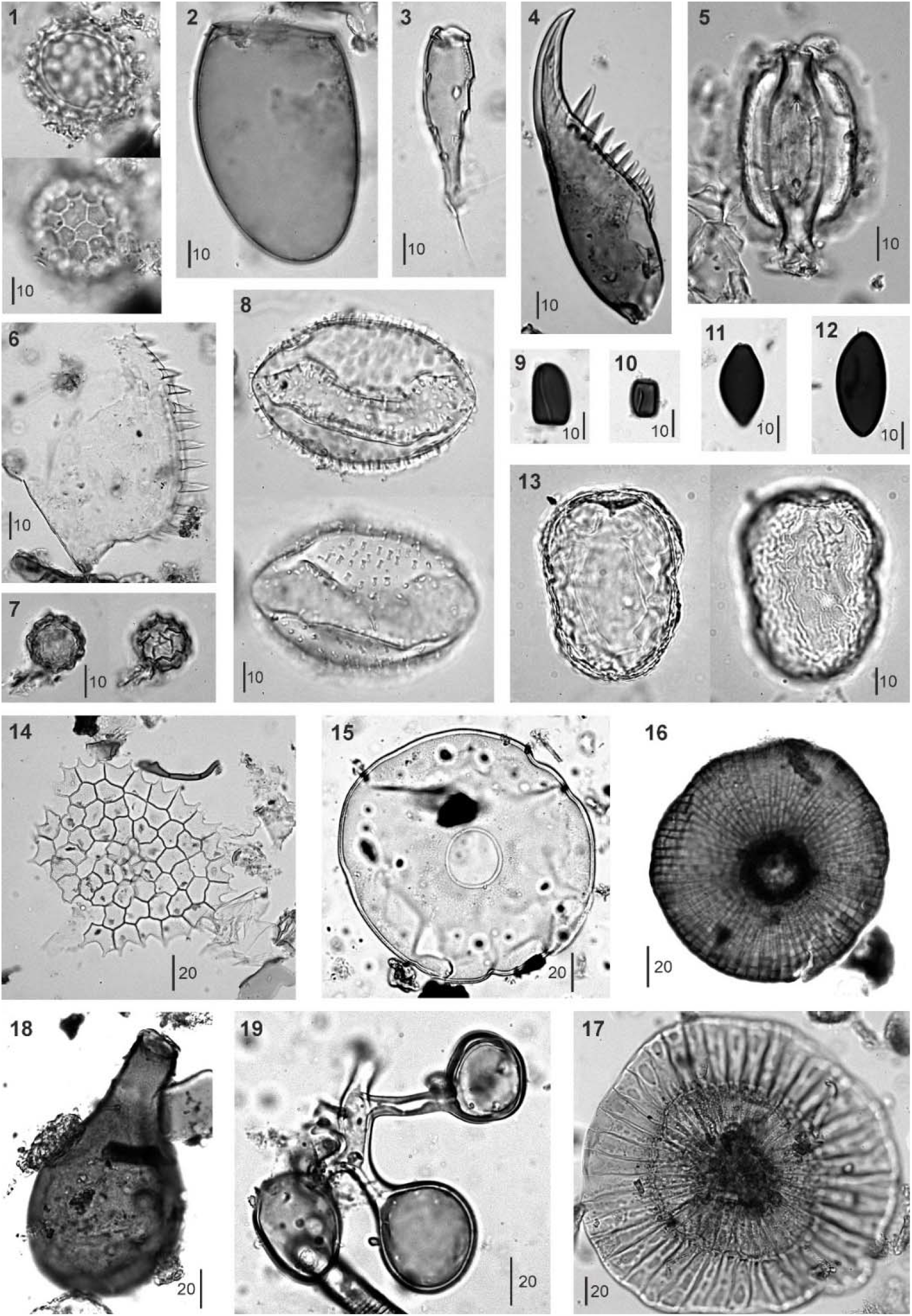
The main NPP (non-pollen palynomorphs) identified in Sant Maursici core SMA13-4A-1G. 1, NPP16; 2, *Neorhabdocoella* egg (tardigrada); 3, Acari leg; 4, Chironomid mandible; 5, *Pinus* stoma; 6, Cladocera mandible; 7, NPP7; 8, NPP14; 9-10, *Sporormiella* spores (fungi); 11, *Sordaria* spore (fungi); 12, *Podospora* spore (fungi); 13, NPP18; 14, Pediastrum coenobium (alga) 15, *Arcella* shell (tecamoebae) 16-17, *Microthyrium* ascomata (fungi); 18, *Habrotrocha* shell (rotifera); 19, *Glomus* spores (fungi). Bar sizes in μm.

A review of the pollen diagram, whether considering the main or the reduced pollen sum, quickly reveals a remarkable constancy in vegetation composition over time (Fig. 3). Indeed, conifer forests, predominantly composed of *Pinus*, have extensively covered the catchment throughout the 20th century. Similarly stable were the abundances of Abies, the second most significant tree species in these forests, and *Rhododendron*, the predominant understory shrub, represented in the diagram by Ericaceae pollen. This constancy also extends to trees from lower elevations, primarily deciduous species from the montane forest belt (notably *Betula, Corylus*, and deciduous *Quercus*), as well as lowland species from Mediterranean sclerophyll forests, such as evergreen *Quercus* and *Olea*. The primary observable changes were in plant cover, as suggested by pollen accumulation rates (PAR), and fire regime, as indicated by charcoal influx. PAR values, largely influenced by arboreal pollen, serve as a reliable proxy for total forest cover. Notably, almost all charcoal particles were smaller than 150 μm, making them effective indicators of regional fire trends.

The diagram starts in the early 1910s, coinciding with the onset of hydroelectricity exploitation within the PNAESM, a period marked by a minor increase in forest abundance and cover, which stabilized around 1940, just before the hydroelectric industry reached its peak development. This rise in conifer forests was accompanied by a slight decline in other forest elements, notably *Betula*, known for successfully colonizing forest clearings in the subalpine belt (Dubois et al., 2020). It is conceivable that the forest recovery, following an initial clearing not captured in the diagram, was facilitated by a reduction in fire incidence, which remained at minimal levels during this time. During this phase, temperatures hovered around the average for the 1930-2020 period, while precipitation was on the rise, peaking in the late 1930s (Fig. 5). This climatic conditions likely supported the observed forest increase. Thus, the modest forest growth noted up until the 1940s may have been the result of a synergistic effect of increasing precipitation and reduced fire incidence. After 1940, forest cover reverted to previous levels, coinciding with a rise in temperatures and a decrease in precipitation.

**Figure 5.**
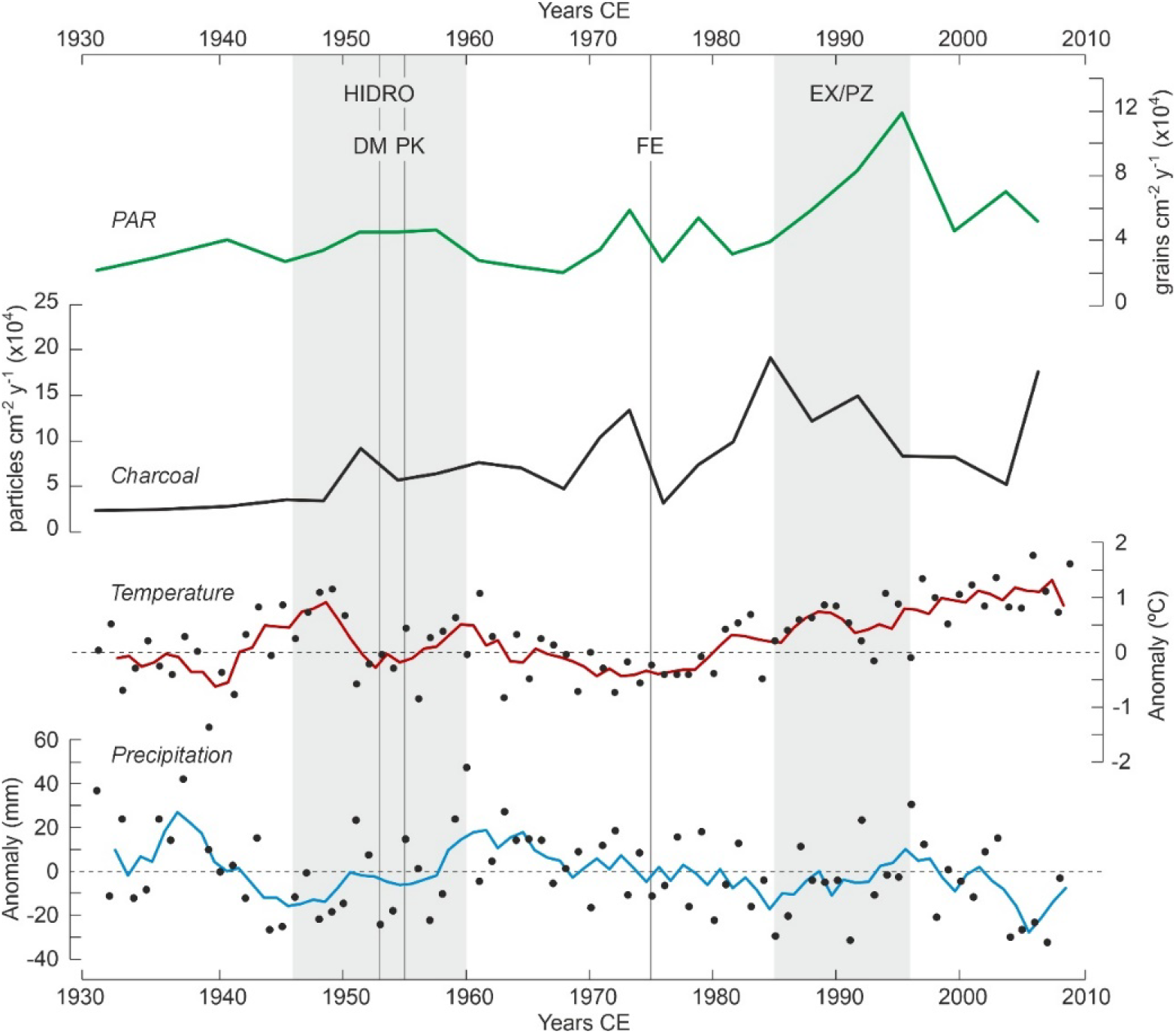
Comparison of pollen accumulation rates (PAR) as a proxy for forest cover, and charcoal influx, as a proxy for fire incidence, with annual average temperature and precipitation trends for the PNAESM. Climatic values are expressed as anomalies and their trends are smoothed using a 4-year moving average (raw data from Sigró et al., 2022).

During the peak period of hydroelectric exploitation (1946-1960), the vegetation composition remained stable, and forests saw a second small recovery, this time solely in terms of plant cover. A notable increase in *Plantago* around the time of the lake damming (1953) likely reflects the transient surge in ruderal vegetation due to local disturbances from the dam construction activities. The establishment of the PNAESM in 1955 did not result in discernible shifts in vegetation composition. Interestingly, an increase in fire incidence coincided with the growth in forest cover, which might be attributed to an increase in available plant fuel for combustion. However, this hypothesis requires further evidence for conclusive validation. These periods of increased plant cover and fire incidence occurred alongside trends of decreasing temperatures and average precipitation, suggesting a minimal climatic influence on these patterns.

After 1960, there was a reduction in plant cover without significant changes in the composition of either forested or non-forested vegetation. This period was characterized by low to intermediate fire incidence and a slight decrease in temperature and precipitation, suggesting that the interplay between climate and fire might have played a significant role. Notably, the period also saw a reduction in coprophilous fungi, particularly *Sporormiella*, indicating a decline in grazing activities. This reduction could reflect a broader decrease in grazing within the PNAESM or result from the local flooding of pastures following dam construction. A third resurgence in forest cover was observed shortly before the imposition of restrictions on forest exploitation in 1975, with no notable changes in community composition. This increase in forest cover once again occurred alongside a significant peak in fire incidence, lending support to the potential link between fire intensity and fuel availability discussed earlier. The lack of significant shifts in temperature and precipitation, which remained stable around the average, bolsters this interpretation.

The imposition of general restrictions on forest exploitation, in 1975, was paralleled by an initial decline in forest cover, followed by an increase and then another decline by 1980. This initial decline occurred alongside a significant reduction in fire incidence, which then surged, reaching its highest levels in the mid-1980s. The fluctuating forest cover values indicate that the forests did not immediately expand in response to the cessation of forest exploitation. It is also possible that the strong fire increase would have contributed to reducing forest cover thus compensating for an eventual forest expansion due to the exploitation cessation. We will come back to this point later. Climate likely played a secondary role in these dynamics, as temperatures were slightly rising and precipitation levels were relatively stable, despite a notable drop coinciding with the peak in fire incidence in the mid-1980s.

During the period of park expansion and the establishment of the peripheral protection zone (1985-1996), forest cover experienced a major increase, reaching its peak in the mid-1990s. The composition of the forest remained constant throughout this time. This increase in forest cover coincided with a notable decline in fire incidence, alongside rising temperatures and increasing precipitation. This pattern suggests that the reduction in fire incidence, combined with climatic conditions conducive to forest growth, supported the expansion of forest cover. At the start of this phase, a simultaneous decline was observed in Pinus stomata and *Microthyrium* ascomata. These NPP exhibited very similar trends throughout the entire diagram, indicating a potential relationship between them. *Microthyrium*, a genus of saprophytic fungi that thrives on leaf litter (Wu et al., 2011), and Pinus stomata, which signal the presence of leaf tissue fragments of local origin in sediments (García Álvarez et al., 2009), can both serve as indicators of sedimentary leaf litter accumulation, particularly from *Pinus*.

Determining the precise causes behind the inferred decline in leaf litter observed in Sant Maurici sediments during the 1980s, and its chronological alignment with the expansion of the park and the creation of the peripheral protection zone, remains challenging given the current evidence. Notably, the surge in *Pinus* stomata immediately following its sharp decline aligns with an uptick in *Plantago* pollen from 1980 onwards. *Plantago* has been identified as an indicator of local disturbance, notably during dam construction activities mentioned earlier. Around this time, park infrastructures underwent renovation and enhancement, with improvements made to access routes, likely contributing to local vegetation disturbances. Consequently, there was a significant rise in visitor numbers, which could account for the sustained increase in *Plantago* presence up to the present. However, there appears to be no direct link between these developments and the significant decline in leaf litter during the 1980s.

A potential explanation for the observed phenomena might relate to the regime of forest exploitation. Logging, particularly for wood extraction, significantly increases the volume of leaf litter on the forest floor by leaving cut branches behind. A portion of this litter eventually makes its way into the lake and its sediments, thereby increasing the content of cuticles where stomata and *Microthyrium* ascomata remain after the decay of more labile organic matter. Consequently, these residues could serve as indicators of forest exploitation. Their continued presence beyond 1975, the year when such practices were officially restricted, suggests that actual cessation of forest exploitation might not have occurred until 1980, coinciding with the implementation of stricter restrictions. This timeline aligns more closely with the amazing increase in forest cover – possibly resulting from either effective expansion of the forested area, an increase in tree density, or both – and the notable decrease in fire incidents starting from 1980.

Following park expansion, there was another decline in forest cover, which occurred alongside a period of low fire incidence (with the exception of the last sample, indicating a recent fire increase), rising temperatures and decreasing precipitation. It is plausible that, with the increased restrictions on human activities, climatic factors became the dominant force. The warming and drying trend observed since the mid-1990s may have contributed to the reduction in forest cover, which attained values similar to the 1970s.

## 5. Discussion and conclusions

### 5.1. Summary of main trends

The above results indicate that the composition of both forested and non-forested vegetation within the Lake Sant Maurici catchment area remained relatively stable throughout the 20th century, demonstrating significant resilience to changes in climate and human activities. This resilience mirrors patterns observed during the Bronze Age and Middle Ages when the catchment area was proposed to have served as a microrefugium for forests amidst widespread deforestation for grazing expansion in the Pyrenean highlands (Rull et al., 2022). However, human impacts during these earlier periods were limited to low-intensity activities such as forest clearing for local cultivation and grazing by nomadic or seasonally migrating societies. In contrast, the 20th century saw a marked intensification of human activities within the PNAESM, including widespread hydroelectric exploitation through lake damming, extensive grazing, forest management for wood extraction, and tourism development, among others. This intensification, coupled with shifts in temperature and precipitation, created a complex scenario of evolving natural and anthropogenic forces and their interactions. Despite these highly dynamic environmental conditions, the vegetation of Sant Maurici still exhibited remarkable resilience in terms of its composition. Climatic and human influences primarily affected plant cover, which was overwhelmingly dominated (90% or more of pollen content) by conifer forests resembling those of the present day, while non-forested communities played a minor role.

The lowest forest cover values were observed from the beginning of hydroelectric exploitation in the 1910s until the late 1970s. During this period, significant events such as the peak of hydroelectric industry activity, the damming of the lake, or the establishment of the national park appeared to exert little or no influence on forest dynamics. Instead, regional fires and climatic trends, along with their potential interactions, played a more pivotal role in the subtle fluctuations of forest cover. It was not until 1980, coinciding with a decline in fire incidence and the expansion of the park alongside stricter protection measures, that forest cover began to significantly increase. Warming trends during this period likely also facilitated forest expansion. In essence, anthropogenic factors emerged as the primary drivers of forest change, with climatic conditions acting as a modulating influence. Interestingly, human interventions such as the lake damming, the establishment of the PNAESM, and the initial restrictions on forest exploitation before 1975 did not reach a threshold of significant impact on forest cover. However, the park’s expansion and the intensification of exploitation restrictions from 1980 onwards surpassed this tipping point. In recent decades, the combination of warming and drying trends has played a crucial role in reducing forest cover, particularly in the absence of logging activities.

### 5.2. Comparisons with other localities and proxies

Given the scarcity of comparable highland palynological records from the last century, the findings of this study can be compared with the mid-elevation Montcortès record and various dendrochronological analyses conducted within the PNAESM.

Lake Montcortès, located 28 km to the south within the lower montane belt, presents a stark contrast in both vegetation and human historical impact compared to Lake Sant Maurici. The regional vegetation around Lake Montcortès is a diverse mosaic comprising deciduous, mixed, and evergreen forests dominated by various oak and pine species, alongside meadows, pastures and cultivated fields (Mercadé et al., 2013). This region has experienced significant human disturbances since Roman times, marked by three major deforestation events that substantially reduced forest cover (Rull & Vegas-Vilarrúbia, 2022). The present-day forests emerged through natural recovery following a 40% reduction in 1800 CE. Throughout the 20th century, these forests showed minimal compositional change, with the only notable shift being a general increase in forest cover, peaking shortly after 1980. Thus, the constancy in composition and variations in forest cover are shared characteristics between the Sant Maurici and Montcortès records. Further studies are required to determine if this pattern represents a broader regional phenomenon and to pinpoint the underlying drivers. A unique difference between the two records is the increase in *Cannabis* pollen at Montcortès during the 1980s, attributed to the extensive cultivation of hemp in the adjacent southeast lowlands (Rull & Vegas-Vilarrúbia, 2023), a feature not observed in the Sant Maurici record. This particular aspect warrants further investigation and will be explored in greater detail in the future through pollen dispersion modeling.

Dendrochronological studies in the PNAESM and its surrounding areas have primarily focused on reconstructing past climates (e.g., Büntgen et al., 2017), documenting snow avalanches (e.g., Muntán et al., 2012), and analyzing tree-growth responses, particularly in *Pinus uncinata*, to environmental drivers, with a notable emphasis on climatic conditions. This discussion will concentrate on the latter category of studies, especially those that can enhance our understanding of the paleoecological reconstruction of Sant Maurici. Over time, the growth of P. *uncinata* trees has been positively correlated with temperatures during the growing season, which includes spring and summer (Hevia et al., 2018). This correlation may help clarify some of the interpretations related to temperature increases and forest cover expansion observed in the study. However, responses to climatic conditions can vary among pine populations. For instance, black pine trees at lower elevations within the park are more adversely affected by summer drought conditions than those at higher elevations (Galván et al., 2014). Given that Sant Maurici is situated at a higher elevation, the impact of summer drought on its forests is expected to be less significant compared to lower elevation areas. Consequently, temperature emerges as the primary factor influencing forest growth in this context, which supports previous interpretations.

Other factors, such as the duration of the winter snow season and the depth of the spring snowpack, also play a significant role in affecting pine growth. Indeed, longer snow seasons and deeper snowpacks have been shown to negatively impact tree growth, irrespective of the atmospheric temperatures during the growing seasons (Sanmiguel-Vallelado et al., 2019). Consequently, warmer conditions, which lead to shorter winters and shallower spring snowpacks, could potentially benefit the growth of black pine trees. Once more, temperature emerges as a fundamental driver for forest growth.

Recent dendroecological surveys have also demonstrated an increase in forest dieback on the Pyrenees linked to current warming and drought intensification (Marqués et al., 2022; Valeriano et al., 2023). This is a worldwide phenomenon that has intensified in the last decades (Anderegg et al., 2013) and is clearly visible in the PNAESM and its surroundings (Fig. 6). This massive forest dieback could help explain the reduction in forest cover observed in the Sant Maurici record starting at the mid-1990s and linked to warming and drying trends.

**Figure 6.**
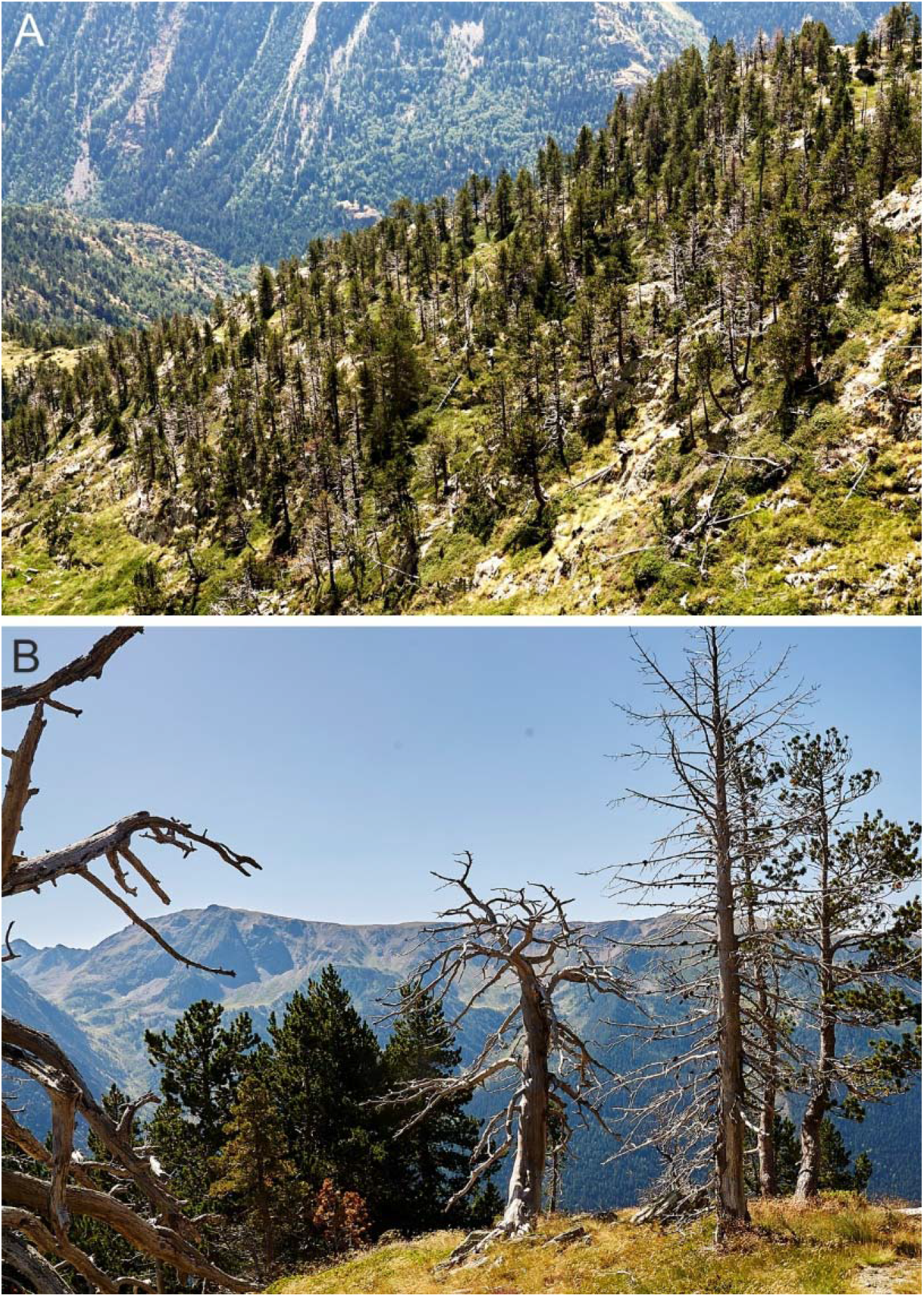
Black pine (*P. mugo subsp. uncinata*) dieback in the surroundings of Lake Naorte, situated 27 km NE from Lake Sant Maurici. A) Black pine forest with a large proportion of dead trees. B) Close-up of the same forest patch showing some dead pines and others alive with signs of drying (brown tones). Photos: V. Rull.

### 5.3. Conservation insights

The high compositional resilience of the forests in Sant Maurici, coupled with their sensitivity to changes in forest cover due to environmental shifts, may have significant implications for conservation. The continuation of the warming and drying trends predicted by the IPCC for this century (Masson-Delmotte et al., 2021) is likely to exacerbate the reduction of forest cover. Based on the results presented, substantial compositional changes are not anticipated unless temperature and/or precipitation reach physiological thresholds critical to tree species with varying climatic tolerances.

A recent ecophysiological study in the PNAESM indicated that *Pinus uncinata* exhibits high tolerance to variations in precipitation, while *Betula pendula* is particularly vulnerable to low precipitation levels, and *Rhododendron ferrugineum* is adversely affected by brief drought events (Fernàndez-Martínez et al., 2016). Consequently, continued drying and a potential increase in drought frequency could further solidify the dominance of black pine. Furthermore, if the anticipated reduction in forest cover leads to a decrease in tree density (Fig. 6), R. *ferrugineum*, which thrives in the shade beneath black pine canopies, might be replaced by shrubs more resistant to sunlight exposure (Fernàndez-Martínez et al., 2016). Additionally, B. *pendula*, frequently found colonizing high-mountain forest clearings, might see an expansion in its populations as long as its moisture requirements are met. This suggests that conservation strategies should prioritize understanding the species-specific responses to potential warming and drying trends, rather than focusing solely on the forest as a whole.

A topic of considerable debate is whether ecosystems should be preserved in their current state or restored to a condition resembling their pre-anthropic state, as inferred by paleoecological records (Willis et al., 2010). For Sant Maurici, the earliest paleoecological records available date back to the Bronze Age (ca. 4.3 to 3.5 cal kyr BP). These records show that the forest composition was remarkably similar to today and that human impact at that time was minimal (Rull et al., 2021). This similarity indicates that the contemporary forest state closely mirrors its natural condition, suggesting that restoration efforts might not be as critical here as in other Pyrenean highlands. Historically, many of these areas were forested but have since been transformed into extensive pastures due to human activities (García-Ruiz et al., 2020; Rull & Vegas-Vilarrúbia, 2021).

Special attention must be given to fire incidence. The forest dieback observed over recent decades not only diminishes forest cover and impacts shadow-loving understory shrubs but also leads to a significant increase in highly flammable deadwood (Fig. 6). The sharp rise in fire incidence seen in the early 2000s could stem from the accumulation of dead trees. Should the current trend of warming and drying persist, fires could pose a major threat to the forests within the PNAESM.

The findings of this study highlight that the most impactful conservation strategies developed to date included establishing a peripheral buffer zone and implementing a strict ban on forest exploitation. Consequently, further expansion of the park, adhering to these same regulations, would likely be advantageous for conservation efforts. If such expansion encompasses areas that were once forested but have since been converted into pastures, restoration initiatives should be seriously considered. For these areas, conducting new paleoecological studies tailored to each specific catchment area would be essential. As it occurred with the creation of the PNAESM and the progress of conservation regulations, an eventual expansion of the park would enhance conflicts with private and communal owners (Gil-Farrero, 2021, 2022). Therefore, further negotiations among all the actors involved would be necessary.

## Acknowledgments

This work was funded by the Autonomous Organism of National Parks, projects RECREO (OAPN-387/2011) and LACEN (OAPN-24505/2017), the AGAUR project GREB (2021SGR00315) and the CERCA Programme, Generalitat de Catalunya. Fieldwork was completed with the collaboration of Fernando Barreiro, Miguel Bartolomé, Matias Frugone, Sandra Garcés-Pastor, Maria Leunda, Julià López, Carlos Pérez, Pedro Sánchez and Blas Valero-Garcés. Fieldwork permits were provided by the National Park “Aiguestortes i Estany de Sant Maurici”.

## References

Anderegg, W.R.L., Kane, J.K., Anderegg, L.D.L. 2013. Consequences of widespread tree mortality triggered by drought and temperature stress. Nature Climate Change 3, 30–36.

Appleby P.G., Oldfield F. 1983. The assessment of 210Pb data from sites with varying sediment accumulation rates. Hydrobiologia 103, 29–35.

Baker, A.G., Bhagwat, S.A., Willis, K.J. 2013. Do dung fungal spores make a good proxy for past distribution of large herbivores? Quaternary Science Reviews 62, 21–31.

Bennett, K.D., Willis, K.D. 2001. Pollen. In: Smol, J.P., Birks, H.J.B., Last, W.M. (eds.), Tracking Environmental Change Using Lake Sediments. Volume 3: Terrestrial, Algal and Siliceous Indicators. Kluwer, New York, pp. 5–32.

Büntgen, U., Krusic, P.J., Verstege, A., Sangüesa-Barreda, G., Wagner, S., Camarero, J.J., Ljungqvist, F.C., Zorita, E., Oppenheimer, C., Konter, O., Tegel, W., Gärtner, H., Cherubini, P., Reinig, F., Esper, J. 2017. New tree-ring evidence from the Pyrenees reveals western Mediterranean climate variability since Medieval times. Journal of Climate 2017, 30, 5295–5318.

Calero, M.A., Valero-Garcés, B., Rull, V., Vegas-Vilarrúbia, T., Garcés-Pastor, S., López-Vila, J., Camarero, J.J. 2016. El registro sedimentario del lago Sant Maurici (Pirineos Centrales). Geogaceta 59, 11–14.

Calero, M.A., Schulte, L., Rull, V., Vegas-Vilarrúbia, T. 2022. Profunditat de l’Estany de Sant Maurici abans de l’any 1953. XII Jornades de Recerca al Parc Nacional d’Aigüestortes i Estany de Sant Maurici, Espot, pp. 53–56.

Camarero, J.J., Garcés-Pastor, S., Gutiérrez, E., Rull, V., Vegas-Vilarrúbia, T., Cañellas-Boltà, N., Sangüesa, G., Trapote, M.C., Clavaguera, A., Calero, M.A., Galván, J.D., Sánchez, R., Giralt, S., Valero-Garcés, B. 2016. Reconstruyendo la historia de los bosques pirenaicos. In: Proyectos de Investigación en Parques Nacionales (2011-2014). O.A. Parques Nacionales, Madrid, pp. 141–156.

Carrillo, E., Aniz, M. 2013. Guía del Parque Nacional de Aigüestortes i Estany de Sant Maurici. O. A. Parques nacionales & Generalitat de Catalunya, Madrid.

Catalan, J., Vilalta, R., Weitzmann, B., Pigem, C., Ventura, M., Aranda, R., Comas, E. 1997. L’Obra Hidràulica en les Pirineus. Avaluació, Correcció i Prevenció de l’Impacte Mediambiental. ENHER-Fundació La Caixa-FECSA, Barcelona.

Catalan, J., Pelachs, A., Gassiot, E. et al. 2013. Interaccion entre clima y ocupacion humana en la configuracion del paisaje vegetal del Parque Nacional de Aiguestortes i Estany de Sant Maurici a lo largo de los ultimos 15.000 anos. In: Ramirez L and Asensio B (eds) Proyectos de Investigación en Parques Nacionales: 2009-2012. Madrid: Organismo Autonomo de Parques Nacionales, pp.71–92.

Cugny, C., Mazier, F., Galop, D. 2010. Modern and fossil non-pollen palynomorphs from the Basque mountains (western Pyrenees, France): the use of coprophilous fungi to reconstruct pastoral activity. Vegetation History and Archaeobotany 19, 391–408.

Davies, A.L., Colombo, S., Nick, H. 2014. Improving the application of long-term ecology in conservation and land management. Journal of Applied Ecology 51, 63–70.

Dubois. H., Verkasalo, E., Claessens, H. 2020. Potential of birch (Betula pendula Roth and B. pubescescens Ehrh.) for forestry and forest-based industry sector with the changing climatic and socio-economic context of western Europe. Forests 11, 336.

Esteban, A. 2003. La humanización de las altas cuencas de la Garona y las Nogueras (4500 aC – 1955 dC). Ministerio de Medio Ambiente, Madrid.

Fernàndez-Martínez, J., Alba Fransi, M., Fleck, I. 2016. Ecolophysiological responses of Betula pendula, Pinus uncinata and Rhododendron ferrugineum in the catalan Pyrenees to low summer rainfall. Tree Physiology 36, 1520–1535.

Galván, J.D., Camarero, J.J., Gutiérrez, E. 2014. Seeing the trees for the forest: drivers of individual growth responses to climate in Pinus uncinata mountain forests. Journal of Ecology 102, 1244–1257.

García Álvarez, S., Morla Juaristi, C., Solana Gutiérrez, J., García-Amorena, I. 2009. Taxonomic differences between Pinus sylvestris and P. uncinata revealed in the stomata and cutilce characters for use in the study of fossil material. Review of Palaeobotany and Palynology 155, 61–68.

García-Ruiz, J.M., Tomás-Faci, G., Diarte-Blasco, P., Montes, L., Domingo, R., Sebastián, M., Lasanta, T., González-Sampériz, P., López-Moreno, J.I., Arnáez, J., Beguería, S. 2020. Transhumance and long-term deforestation in the subalpine belt of the Central Spanish Pyrenees: an interdisciplinary approach. Catena 195, 104744.

Gauthier, E., Jouffroy-Bapicot, I. 2021. Detecting human impacts: non-pollen palynomorphs as proxies for human impact on the environment. In: Marret, F., O’Keefe, J., Osterloff, P., Pound, M., Shumilovkikh, L. (eds.), Applications of Non-Pollen Palynomorphs: from Palaeoenvironmental Reconstruction to Biostratigraphy. Geological Society Special Publications 511, London, pp. 233–244.

Gil-Farrero, J. 2021. Un segle de canvis en la gestió dels recursos naturals al Parc Nacional d’Aigüestortes i Estany de Sant Maurici. XII Jornades sobre recerca al Parc Nacional d’Aigüestortes i Estany de Sant Maurici, Espot, pp. 207–216.

Gil-Farrero, J. 2022. Conservación, divulgación e imagen pública de la naturaleza durante el franquismo; El Parque Nacional de Aigüestortes i Estany de Sant Maurici. Rubrica Contemporanea 11, 249.

Hevia, A., Sánchez-Salguero, R., Camarero, J.J., Buras, A., Sangüesa-Barreda, G., Galván, J.D., Gutiérrez, E. 2018. Towards a better understanding of long-term wood-chemistry variations in old-growth forests: a case study on ancient Pinus uncinata trees from the Pyrenees. Science of the Total Environment 625, 220–232.

Lee, C.M., van Geel, B., Gosling, W.D. 2022. On the use of spores of coprophilous fungi preserved in sediments to indicate past herbivore presence. Quaternary 5, 30.

López-Vila, J., Montoya, E., Cañellas-Boltà, N., Rull, V. 2014. Modern non-pollen palynomorphs sedimentation along an elevational transect in the south-central Pyrenees (southwestern Europe) as a tool for Holocene paleoecological reconstruction. Holocene 24, 327–345.

Marqués, L., Ogle, K., Peltier, D.M.P., Camarero, J.J. 2022. Altered climate memory characterizes tree growth during forest dieback. Agricultural and Forest Meteorology 314, 108787.

Masson-Delmotte, V., P. Zhai, A. Pirani, S.L. Connors, C. Péan, S. Berger, N. Caud, Y. Chen, L. Goldfarb, M.I. Gomis, M. Huang, K. Leitzell, E. Lonnoy, J.B.R., Matthews, T.K. Maycock, T. Waterfield, O. Yelekçi, R. Yu, and B. Zhou. 2021. Climate Change 2021: The Physical Science Basis. Cambridge University Press, Cambridge.

Mercadé, A., Vigo, J., Rull, V., Vegas-Vilarrúbia, T., Garcés, S., Lara, A., Cañellas-Boltà, N. 2013. Vegetation and landscape around Lake Montcortès (Catalan pre-Pyrenees) as a tool for palaeoecological studies of lake sediments. Collectanea Botanica 32, 87–101.

Moore, P.D., Webb, J.A., Collinson, M.E. 1991. Pollen Analysis. Blackwell, Oxford.

Muntán, E., García, C., Oller, P., Martí, G., García, A., Gutiérrez, E. 2009. Reconstructing snow avalanches in the Southeastern Pyrenees. Natural Hazards and earth System Science 9, 1599–1612.

Reille, M. 1992. Pollen et Spores d’Europe et d’Afrique du Nord. URA-CNRS, Marseille.

Rodríguez, J.-M., Pérez-Obiol, R., Pérez-Haase, A., Nadal-Tersa, J., Sánchez-Morales, M., Cunill-Artigas, R., Pèlachs, A. 2021. Recerca paleoambiental a l’Estany de la Bassa. XII Jornades de Recerca al Parc Nacional d’Aigüestortes i Estany de Sant Maurici, Espot, pp. 153–162.

Rodríguez-Fernández, R. 2010. Parque Nacional de Aigüestortes I Estany de Sant Maurici. Guía Geológica. Ed. Everest, León.

Rull, V. 1987. A note on pollen counting in palaeoecology. Pollen Spores 29, 471–480.

Rull, V. 2014. Time continuum and true long-term ecology: from theory to practice. Frontiers in Ecology and Evolution 2, 75.

Rull, V. 2023. Anticipation, discovery and serendipity in Quaternary paleoecology: personal experiences from the Iberian Pyrenees. Quaternary 6, 42.

Rull, V., Vegas-Vilarrúbia, T. 2011. What is long term in ecology? Trends in Ecology and Evolution 26, 3–4.

Rull, V., Vegas-Vilarrúbia, T. 2021. A spatiotemporal gradient in the anthropization of Pyrenean landscapes. Preliminary report. Quaternary Science Reviews 258, 106909.

Rull, V., Vegas-Vilarrúbia, T. 2022. Resilience of Pyrenean forests after recurrent historical deforestations. Forests 14, 567.

Rull, V., Vegas-Vilarrúbia, T. 2023. A recent Cannabis pollen increase on the Iberian Pyrenees. Science of the Total Environment 886, 163947.

Rull, V., Cañellas-Boltà, N., Vegas-Vilarrúbia, T. 2021. Late Holocene forest resilience in the central Pyrenean highlands as deduced from pollen analysis of Lake Sant Maurici sediments. Holocene 31, 1797–1803.

Rull, V., Vegas-Vilarrúbia, T., Corella, J.P., Trapote, M.C., Montoya, E., Valero-Garcés, B. 2021. A unique Pyrenean varved record provides a detailed reconstruction of Mediterranean vegetation and land-use dynamics over the last three millennia. Quaternary Science Reviews 268, 107128.

Sánchez-Cabeza J.A., Masqué, P., Ani-Ragolta, I. 1998. 210Pb and 210Po analysis in sediments and soils by microwave acid digestion. Journal of Radioanalytical and Nuclear Chemistry 227, 19–22.

Sanmiguel-Vallelado, A., Camarero, J.J., Gazol, A., Morán-Tejeda, E., Sangüesa-Barreda, M., Alonso-González, E., Gutiérrez, E., Alla, A.Q., Galván, J.D., López-Moreno, J.I. Detecting snow-related signals in radial growth of Pinus uncinata mountain forests. Dendrochronologia 57, 125622.

Seddon, A.W.R., Mackay, A.W., Baker, A.G., Birks, H.J.B., Breman, E., Buck, C., Ellis, E.C., Froyd, C., Gill, J.L., Gilson, L., et al. (2014) Looking forward through the past: identification of 50 priority research questions in paleoecology. Journal of Ecology 102, 256–267.

Sigró, J., Pérez-Luque, A.J., Pérez-Martínez, C., Vegas-Vilarrubia, T., Esteban-Parra, M. J. 2022. Trends in temperature and precipitation in high mountain areas in Spain from the Spanish Hig Mountain Climate Database. EGU General Assembly, Vienna, Austria, 23–27 May 2022.

Stockmarr, J. 1971. Tablets with spores used in absolute pollen analysis. Pollen Spores 13, 615–621.

Theuerkauf, M., Couwenberg, J. 2018. ROPES reveals past land cover and PPs from single pollen records. Frontiers in Earth Science 6, 14.

Valeriano, C., Tumajer, J., Cazol, A., González, E., Sánches-Salguero, R., Colangelo, M., Linares, J.C., Valor, T., Sangüesa-Barreda, G., Camarero, J.J. 2023. Delineating vulnerability to drought using a process-based growth model in Pyrenean silver fir forests. Forest Ecology and Management 541, 121069.

Vegas-Vilarrúbia, T., Rull, V., Montoya, E., Safont, E. 2011. Quaternary palaeoecology and nature conservation: a general review with examples from the neotropics. Quaternary Science Reviews 30, 2361–2388.

Vigo, J. 2008. L’alta muntanya catalana. Flora i vegetació. Centre Excursionista de Catalunya-Institut d’Estudis Catalans, Barcelona.

Whitlock, C., Larsen, C. 2001. Charcoal as a fire proxy. In: Smol, J.P., Birks, H.J.B., Last, W.L. (eds.), Tracking Environmental Change Using Lake Sediments. Volume 3: Terrestrial, Algal and Siliceous Indicators. Kluwer, New York, pp. 75–97.

Willis, K.J., Birks, H.J.B. 2006. What is natural? The need for a long-term perspective in biodiversity conservation. Science 314, 1261–1265.

Willis, K.J., Araújo, M.B., Bennett, K.D., Figueroa-Rangel, B., Froyd, C.A., Myesr, N. 2007. How can a knowledge of the past help to conserve the future? Biodiversity conservation and the relevance of long-term ecological studies. Philosophical Transactions of the Royal Society B 362, 175–187.

Willis, K.J., Bailey, R.M., Bhagwat, S.A., Birks, H.J.B. 2010. Biodiversity baselines, thresholds and resilience: testing predictions and assumptions using paleoecological data. Trends in Ecology and Evolution 25, 583–591.

Wu, H.X., Schoch, C.L., Boonmmee, S., Bahkali, A.H., Chomnuti, P., Hyde, K.D. 2011. A reappraisal of Microthyriaceae. Fungal Diversity 1, 189–248.

